# Optical Coherence Tomography with Fluorescein Optical Clearing for Transscleral Image Guidance

**DOI:** 10.1101/2025.07.01.661162

**Authors:** Robert Trout, Amit Narawane, Christian Viehland, Vahid Ownagh, Mark Draelos, Al-Hafeez Dhalla, Anthony N. Kuo, Cynthia A. Toth

## Abstract

Optical coherence tomography (OCT) is a non-invasive, three-dimensional imaging modality demonstrated to yield a wealth of high-resolution structural and functional information in a variety of biomedical imaging applications. However, there is a significant limitation on the penetration depth of the modality in most tissues due to their high optical scattering, hindering imaging of more deeply embedded targets. In ophthalmic applications, this makes transscleral imaging of targets such as suprachoroidal injection volumes, lesions, and malignancies challenging. Here, we investigate the novel application of fluorescein, a commonly applied biomedical dye, as an optical clearing agent for increasing the depth of OCT imaging in scleral tissues. The effect is characterized in ex-vivo models including the sclera of porcine and human eyes, examining its time and concentration dependence, reversibility, and potential for application in the enhancement of intrasurgical trabeculectomy image guidance. We demonstrate that fluorescein can serve not only as a biomedical fluorophore, but also as an optical clearing agent capable of significantly increasing OCT imaging depth in tissue. As a common and accessible ophthalmic agent with a long history of clinical application, we believe that this new potential use could have substantial positive impact in the enhancement of biomedical imaging in scattering tissue.

## 2. Introduction

Optical coherence tomography (OCT) is a volumetric imaging modality demonstrated to be a powerful tool in a variety of clinical applications, providing image-based diagnostics and guidance in areas spanning ophthalmology, neurology, oncology, angiology, gynecology, dermatology, and gastroenterology^1^. While these applications yield a wealth of high-resolution, three-dimensional structural and functional information, there is a severe limit on the penetration depth of the modality in most tissues. This can hinder the imaging of targets in many contexts, for example, in ophthalmology, transscleral OCT imaging could prove useful in several applications in the anterior segment, such as angle measurements for glaucoma, full-thickness imaging of malignancy margins, verification of implants and injections, and localization of muscle insertions for strabismus operations^2^. However, the imaging depth in sclera is quite shallow, limiting its potential in these applications. The chief mechanism responsible for this limitation is the high degree of optical scattering exhibited by the sclera (and most other biological tissues) due to its composition of protein and fat structures suspended in a watery medium. As a result, heavy scattering occurs for incident light due to the difference in refractive index between these structures and the surrounding water^3^. In the case of OCT, the portion of light which undergoes extensive scattering as it travels through the tissue is rejected due to the confocal and coherence gating of the modality. While this part of the signal has generally lost any structural information and is analogous to noise, the fact remains that as the image depth increases, more and more of the structural information of the tissue reflectance is lost to the increasing scattering encountered along longer depths of propagation. These losses manifest in an attenuation of OCT signal with image depth, limiting OCT imaging penetration in tissue to 1-2mm in most applications^4,5^.

A substantial amount of work has been dedicated to addressing this limitation in OCT and other imaging modalities, with methods spanning a variety of optical and chemical approaches^6,7^. For in-vivo applications, a promising approach has been found in the application of various biocompatible optical clearing agents, including sugar solutions, glycerol, propylene glycol, and dimethylsulfoxide (DMSO)^8–11^. Diffusion of these agents into the tissue results in a homogenization of its refractive index, reducing (clearing) the optical scattering and providing increased imaging depth in many optical imaging applications including OCT. However, the efficiency of refractive index change for these agents is poor, requiring high concentrations within the tissue to achieve substantial clearing.

More recently, absorbing molecules such as the food dye tartrazine have been demonstrated to be a far more efficient at optical clearing in tissue^12^. These agents paradoxically increase clearing of tissues by virtue of their absorbance properties, due to the coupling of absorbance and dispersion as described by the Kramers-Kronig relations^13^. In this way, peaks in the absorption spectrum are accompanied by increased refractive index at longer wavelengths. If the imaging wavelength is longer than the absorption peak, significant optical clearing can be achieved in tissue with minimal absorption loss, and as the diffusion of the agent into the tissue aqueous raises its refractive index to more closely match that of non-aqueous structures, scattering is reduced. We have previously demonstrated that this mechanism can be leveraged with tartrazine in ophthalmic OCT to enable transscleral imaging^14^, while others have corroborated this in other work with volumetric imaging modalities^15,16^. While promising, given the lack of data regarding safety of its topical use, tartrazine has limited immediate utility for *in vivo* optical clearing.

However, many absorbing dyes beyond tartrazine exist which are commonplace in clinical ophthalmic applications. Among these, by far the oldest and most commonly used is fluorescein^17^. While currently used for its fluorescence properties in the staining of ophthalmic pathologies and anatomy, fluorescein also possesses a strong absorbance peak (490nm) significantly below typical NIR OCT imaging wavelengths like that of tartrazine (425nm). With this, we believe fluorescein has potential to serve as a more clinically compatible absorptive optical clearing agent compared to tartrazine or other agents without such an extensive history of clinical use.

For our pilot application, we aim to demonstrate the potential for fluorescein clearing to address some of the challenges in transscleral OCT imaging due to the sclera’s opacity to optical imaging from its high scattering. This could have implications for image guidance and diagnostics in subscleral surgical maneuvers, injections, lesions, and malignancies. Here we choose to investigate its potential for the enhancement of image guidance in ophthalmic surgery. The high degree of precision demanded by these procedures has driven the development of increasingly sophisticated technologies to advance the capabilities of the surgeon. Recently, optical coherence tomography (OCT) has also been demonstrated as a valuable tool in this regard, adding high-resolution volumetric imaging of the surgical field^18^. As a result, many groups, including our own, have developed high speed microscope-integrated OCT (MIOCT) systems to introduce real time OCT image guidance to the surgical theater^19–22^. These systems provide high resolution, depth-resolved OCT imaging of tissue at live video rates to the surgeon, enhancing surgical guidance and decision making in a variety of scenarios in both the anterior and retina. Combined with the improved image depth afforded by the optical clearing of fluorescein, we believe that there is potential for enhanced image guidance by intrasurgical OCT for transscleral procedures like trabeculectomies.

In this work, we demonstrate for the first time: 1) fluorescein can be used as an optical clearing agent in *ex vivo* studies with porcine and human donor eyes; 2) fluorescein is a reversible optical clearing agent; and 3) the potential for deep-tissue visual guidance in trabeculectomies as well as other procedures and diagnostics. We believe this work presents a valuable step towards *in vivo* optical clearing with highly absorbing molecules.

## 3. Methods

To investigate the clearing effect of fluorescein in scleral OCT imaging, mounted *ex vivo* human and porcine eyes were utilized as models of *in vivo* human imaging (Fig. 1). Clearing effects in imaging were studied first as a function of time and concentration in a controlled setting (Fig. 1b), then in a model intrasurgical trabeculectomy scenario (Fig. 1c).

**Figure 1.**
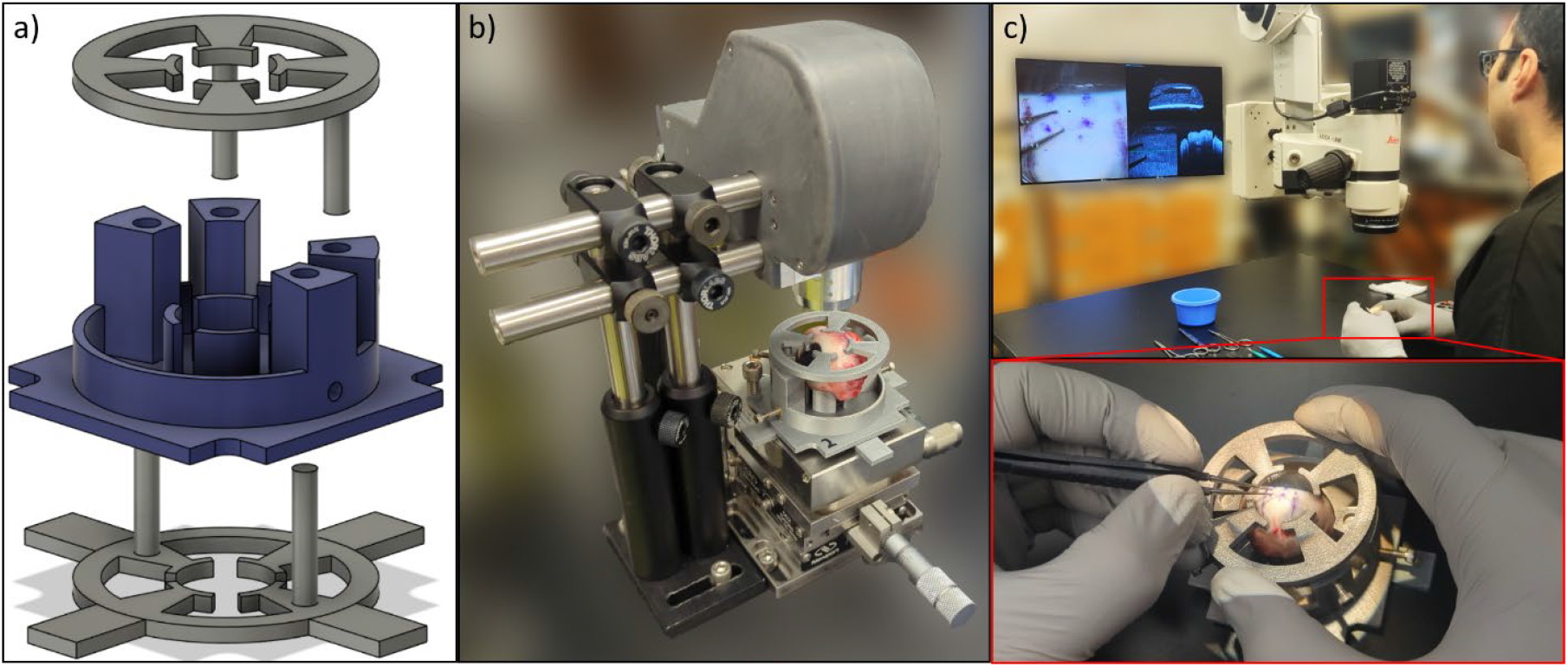
Methods for scleral clearing imaging. a) Mount designed and 3D-printed for repeatable positioning of the ex vivo eyes during imaging. b) Stage-mounted custom anterior-segment OCT scanner setup for time-series imaging. c) Investigational high-speed microscope-integrated OCT for intrasurgical imaging during pilot trabeculectomy surgery (inset).

### 3.1. Sample Preparation

As models of trabeculectomy, all sample eyes had their conjunctiva removed from the area of imaging prior to treatment. To achieve sample stabilization and consistent imaging alignment over time across the *ex vivo* eye samples, a custom mounting solution was designed and 3D printed for repeatable positioning of the eyes relative to the imaging system. This was required for control of imaging variations related to sample positioning, enabling more robust quantitative comparison across eyes. Using rod features and set screws, the height of the sample may be offset from the base while the height of the lid can be adjusted to secure eyes of varying size (Fig. 1a).

After mounting, dosing of eyes was conducted with a sodium fluorescein salt (Aqua Solutions, Deer Park, TX) deionized water solution. To control for drying effects that may occur due to the hyperosmolarity of the fluorescein, an equiosmolar control solution of sodium chloride (non-iodized table salt) was applied for comparison. All eyes were dosed with fluorescein or salt control solution at a rate of 5 drops every 5 minutes over the course of 1 hour.

### 3.2. Time-Series Imaging

To examine the time dependence and repeatability of the clearing effect, control (7% salt) and experimental (30% fluorescein) porcine eyes (n=3) were positioned on an XYZ micrometer stage for aligned imaging using an investigational handheld anterior segment research scanner detailed in previous work (Fig. 1b)^14^. The scan protocol utilized for this study was 1000 a-scans by 128 b-scans averaged 8x over a 13×13mm field of view. Imaging was taken pre-dose, and 10, 20, 40, and 60 minutes timepoints during dosing. This experiment was then repeated with a pair of ex-vivo human donor eyes (n=1) to see how the effect compared in a more realistic model.

The concentration dependence of the clearing effect was characterized by imaging porcine eyes dosed with 2, 5, 10, 15, and 30% fluorescein solutions (n=1 per concentration) over the course of 1 hour.

Reversibility of the clearing effect was characterized by first clearing a porcine eye (n=1) with 30% fluorescein for 1 hour, followed by flushing with 3 drops per second of balanced saline solution (BSS) for 10 minutes, imaged pre-rinse, and after 1, 2, 5, and 10 minutes of flushing.

To make quantitative comparisons across different imaging conditions, a-scans at corresponding positions were selected for each eye. 10 adjacent a-scans were averaged together, and the smoothed result was normalized to the starting pre-dose depth profile of the eye. In this way, the profiles represent change in image signal of each eye relative to the signal in their pre-dose profile, normalizing inter-eye variation that may be present. The distributions of these image depth profiles were then compared to examine the effect of clearing on imaging depth under different conditions.

### 3.3. Intrasurgical Imaging

For intrasurgical imaging (Fig. 1c), an investigational high-speed MIOCT system detailed previously^20^ was used to acquire digital microscopy and OCT scanning in dense/large configuration for snapshots (750 a-scans by 750 b-scans over 15×15mm), and sparse/fast configuration for guidance at a 6Hz volume rate (200 a-scans by 80 b-scans over 8×8mm).

The pilot surgery conducted consisted of a trabeculectomy carried out by an expert surgeon trained in trabeculectomy procedures for glaucoma treatment (Fig. 2). This procedure was selected as a pilot application due to its status as the gold standard treatment for advanced glaucoma, the leading cause of irreversible blindness, and its high degree of surgical precision in the careful placement of incisions, needle insertions, and implants within the eye^23,24^. It is theorized that some of the maneuvers associated with the procedure could potentially benefit from transscleral clearing; with the high scattering of scleral tissue, the surgical target and instrument are often obscured from the surgeon’s sight, leaving the surgeon to rely on memory and indirect cues. Optical clearing of these tissues may aid the visual guidance of the surgery.

**Figure 2.**
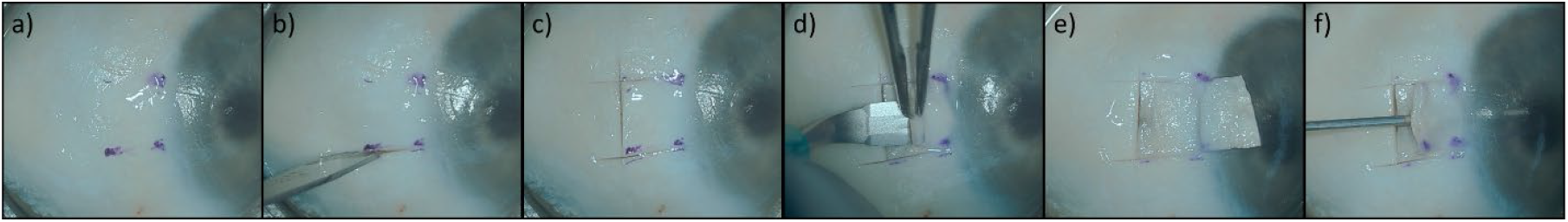
Pilot trabeculectomy procedure for intrasurgical imaging, human ex vivo. Steps of pilot surgery include a-c) Initial incisions, d,e) scleral flap creation, (f) Drainage channel creation.

The pilot procedure was performed *ex vivo* following the removal of the conjunctiva in the target area. Surgical steps included creation of a 4mm sclerotomy flap (Fig. 2a-e), followed by the insertion of a 25G needle at the flap joint into the anterior chamber (AC) of the eye to create a drainage channel (2f).

## 4. Results

### 4.1. Fluorescein Clearing Effect: Porcine *Ex Vivo*

Over the course of 1 hour of dosing porcine eyes with control (7% salt) and clearing (30% fluorescein) solutions (n=3), a noticeable increase in image depth was observed in the sclera between control (Fig. 3a, left) and cleared (right) OCT imaging. For reference, the anterior eye and cornea are positioned on the right side, with the posterior side on the left. All subsequent b-scan imaging in this work will continue to follow this orientation convention. From these profiles, little change in depth of imaging is observed over time in the control case, while in the cleared case there is a reduction in signal higher in the superficial regions of the sclera whose reflectance is reduces due to fluorescein exposure. However, this enables increased penetration depth of unscattered OCT light, improving visibility of deeper scleral layers and subscleral structures including the choroid/ciliary body. While the clearing effect is not sufficient to make out subscleral features on corresponding white-light surgical microscopy (Fig. 3b), plotting the average and standard error of the OCT depth profiles (Fig. 3c), reveals the signal attenuation in the superficial sclera (green arrow) accompanied by significant increases in the OCT signal at greater depths in the bulk sclera (yellow arrow) after dosing for a time period as short as 10 minutes, with the effect appearing to saturate between 40 and 60 minutes. At this point, in clearing, the superficial sclera in the first 0.5mm of depth sees a signal reduction as great as -20dB, while a peak signal gain of approximately +15dB manifests in the scleral tissue at greater depths beyond 1mm. A smaller secondary peak can also be seen near 2.6mm depth as the signal from the choroid/ciliary body increases (red arrow).

**Figure 3.**
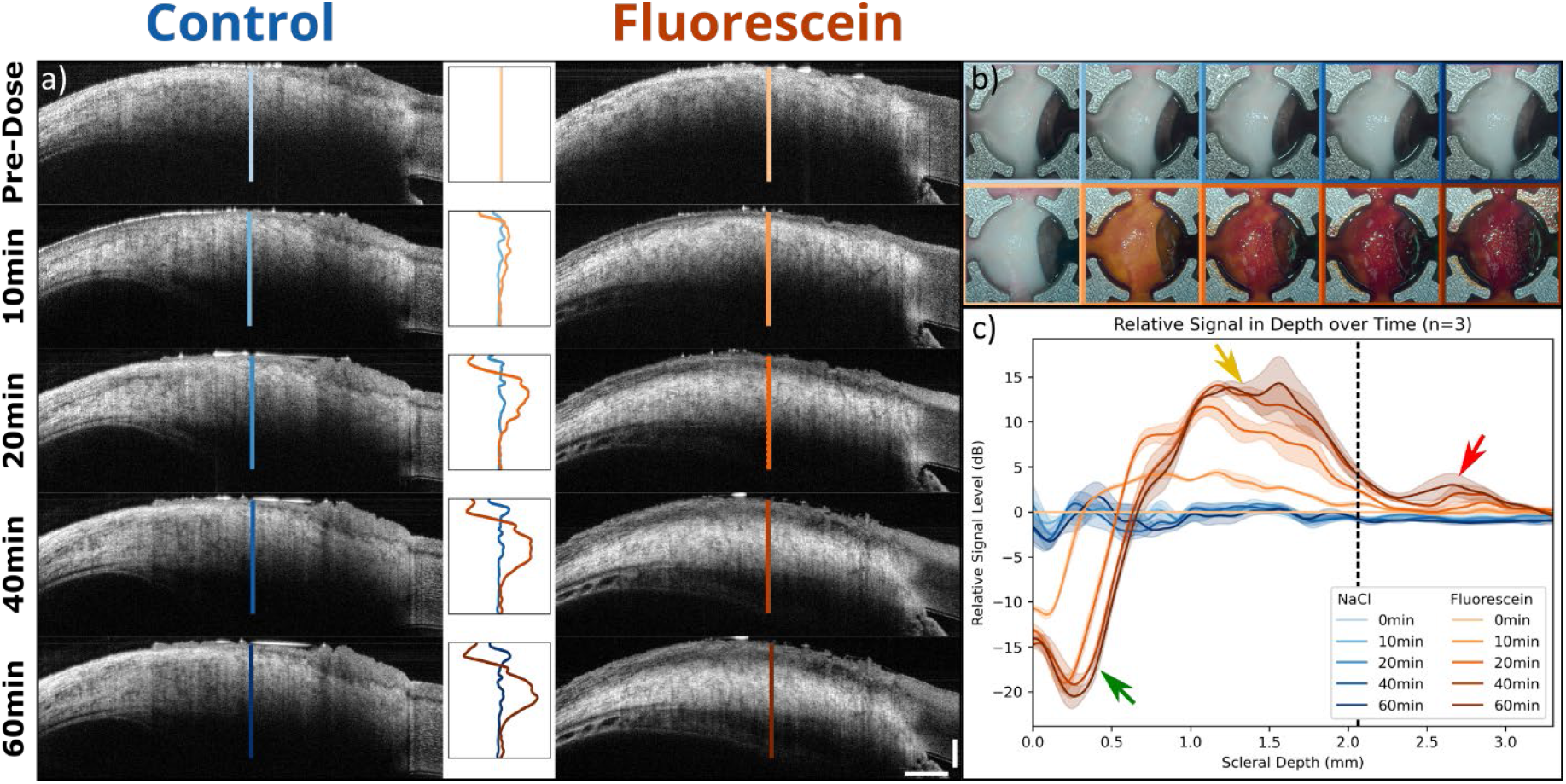
a) Time series OCT imaging of clearing effect over time of 7% salt control solution (left) compared to isoosmolar 30% fluorescein (right) in porcine eyes. Top row is immediately before dosing, followed by 10, 20, 40, and 60 minutes of dose application. Smoothed image signal depth profiles for each image are plotted normalized to the starting profile (middle), with line annotations indicating the position of the plotted a-scan. (b) Corresponding enface surgical microscopy imaging for salt control (top row) and fluorescein (bottom) over time. (c) Average depth profiles across 3 repeats for both conditions, with standard error as shaded curve bounds, with approximate position of superficial sclera (green arrow), bulk sclera (yellow arrow), choroid-scleral junction (dotted black line), and ciliary body/choroid (red arrow). 1mm scale.

The degree of OCT clearing is also dependent on the concentration of the fluorescein solution, as illustrated in Fig. 4 below.

**Figure 4.**
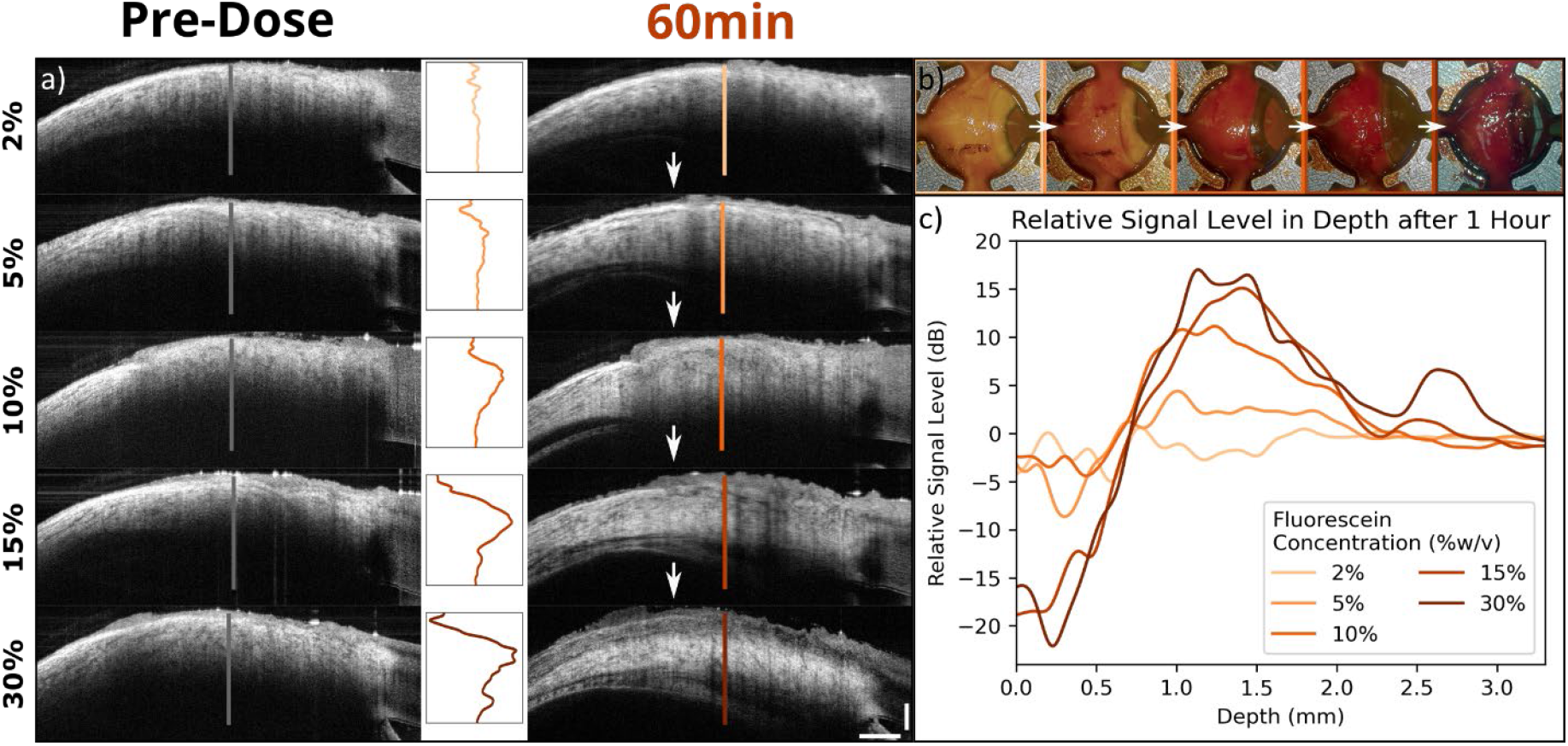
a) OCT imaging of clearing effect before (left) and after (right) 1 hour dosing with fluorescein solutions of varying concentrations. From top to bottom, these concentrations include 2, 5, 10, 15, and 30%. Smoothed image signal depth profiles for each image are plotted normalized to the pre-dose profile (middle), with line annotations indicating the position of the plotted a-scan. (b) Corresponding enface surgical microscopy imaging following dosing at each concentration of fluorescein. (c) Combined plot of profiles across concentrations.

For each eye (n=1), a clearing effect can be visualized between the pre-dose (Fig. 4a, left) and the post 1 hour fluorescein treatment imaging (right). Similar to the previously observed effect of longer dose times, higher concentrations of fluorescein result in the increased clearing of the superficial scleral tissue, reducing OCT signal at the surface while increasing it at greater tissue depths (middle). Plotted together (Fig. 4c), the majority of the effect appears to saturate between 15-30% concentration.

Following clearing treatment with fluorescein, the effect can be readily reversed via flushing of the eye with BSS (n=1). As observed in Fig. 5, after the tissue is cleared with 1 hour treatment with fluorescein (5a, left), flushing of the eye with BSS resulted in a complete reversal of the clearing effect in the OCT imaging after just 10 minutes (right). Thus, the clearing does not appear to be a permanent optical effect on the tissue. The appearance of the eye under white-light microscopy (5b) has not completely returned to its initial appearance following this wash, due to the much greater sensitivity of white-light imaging to lower concentrations of fluorescein. Complete reversal of the yellow shading from this will thus have a much longer timescale on the order of hours compared to OCT imaging. The dynamics of the signal depth distributions are quantified in 5c, where the distribution can be observed to shift deeper relative to the pre-dose distribution (50^th^ percentile annotated with dotted red line) following 1 hour of clearing treatment. This shift then completes a reversal after 5-10 minutes of flushing.

**Figure 5.**
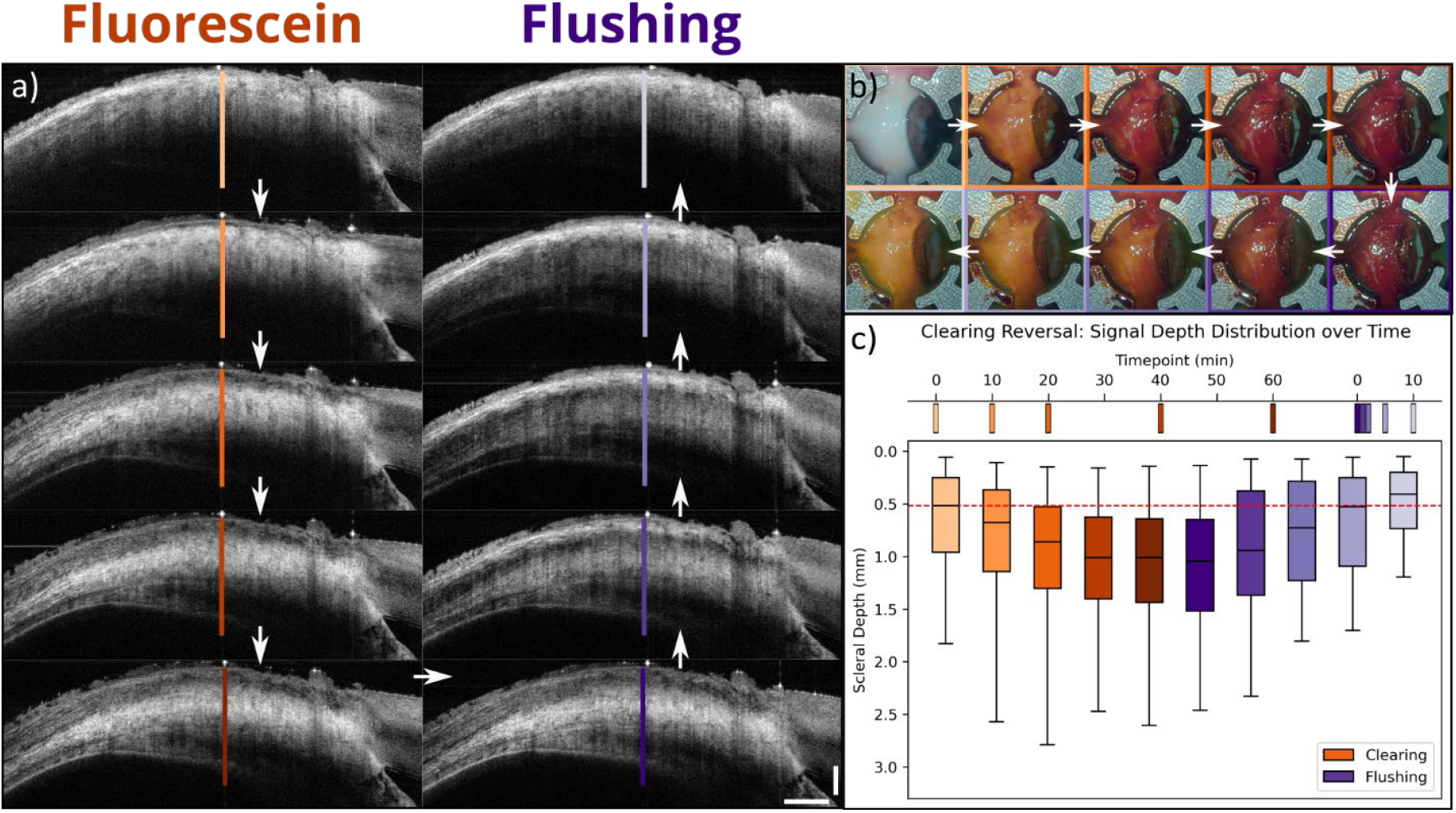
a) Time series OCT imaging of clearing effect over time of fluorescein solution (left) with subsequent reversal of clearing by flushing with balanced saline solution (right). White arrows indicate order in time. Fluorescein dosing image timepoints of (0) pre-dosing followed by 10, 20, 40, and 60 minutes of dosing, then reversal flushing timepoints of 0 (pre-flushing), 1, 2, 5,10 minutes. (b) Corresponding enface surgical microscopy imaging for fluorescein clearing (top row) and reversal flushing (bottom) over time. (c) Imaging timeline (top) and boxplots for signal distribution in depth for each condition and timepoint at locations annotated in (a).

### 4.2. Pilot Trabeculectomy Imaging

Following the 1-hour treatment the trabeculectomy procedure was perform while collecting intraoperative OCT and microscopy imaging. The tissue clearing effect was found to improve OCT visualization of the surgery, particularly in the case of the needle insertion when creating the drainage channel, depicted in Fig. 6.

**Figure 6.**
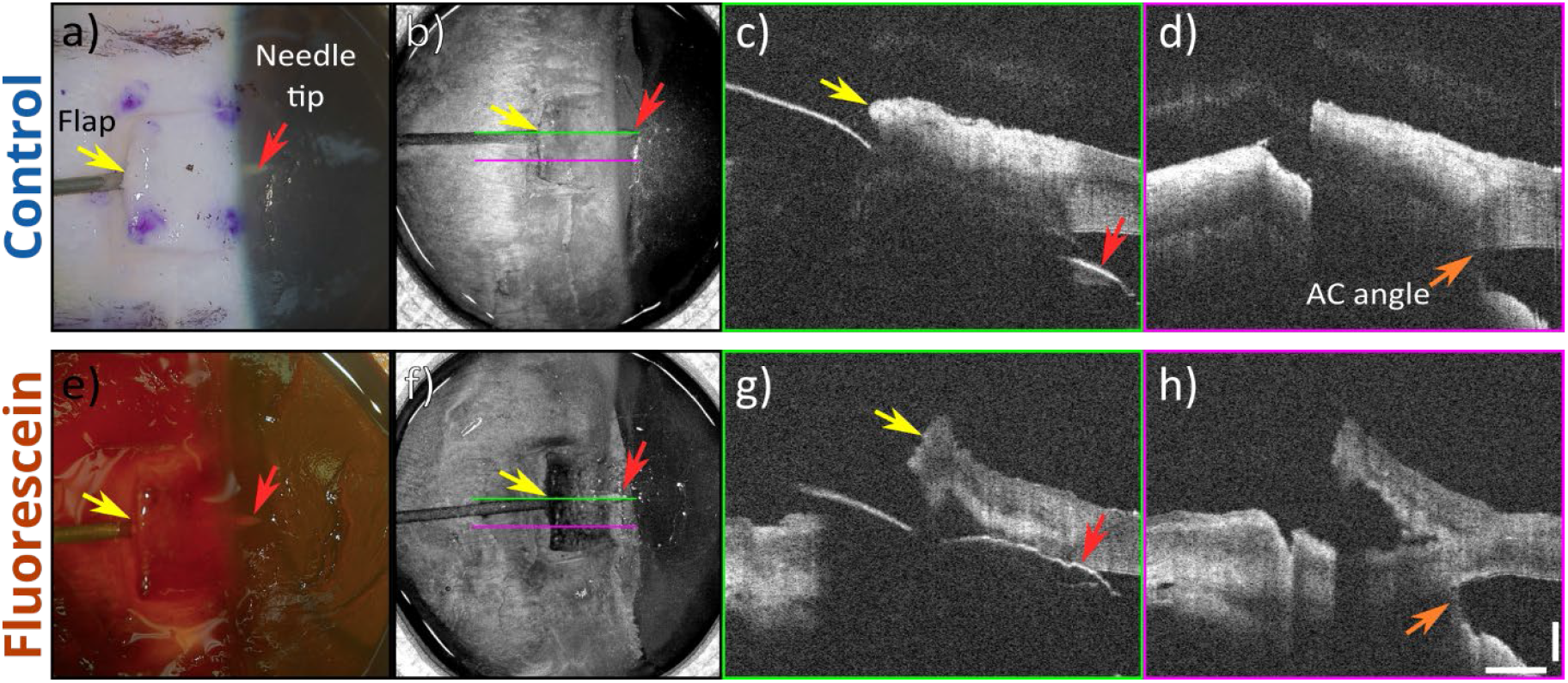
a) Intrasurgical microscopy imaging of ex vivo porcine eye during trabeculectomy channel creation following 1hr exposure to 7% NaCl solution. Scleral flap (yellow arrow) and needle tip (red) visualized. b) Corresponding dense-scan OCT enface max-intensity projection. c) B-scan along needle axis (green); warping in the imaged needle surface is due to optical path length differences from variable flap tissue thickness above the needle. d) B-scan adjacent to needle to visualize features previously shadowed by needle, including the flap bed (purple arrow) and AC angle (orange). e-h) Corresponding 30% fluorescein-cleared imaging. 1mm scale.

While the improvement to the microscopy visualization (a, e) of the needle (red arrow) through the flap (yellow arrow) is limited, the visual obstruction in OCT imaging of the needle (c), and the AC angle (d, orange arrow) is improved dramatically with clearing (g, h), yielding complete visualization of the needle’s length beneath the flap, and the AC angle target for the channel placement.

This enhanced visibility of the needle through the flap enables improved visual tracking of the instrument using high-speed OCT scanning, as demonstrated in Fig. 7 below.

**Figure 7.**
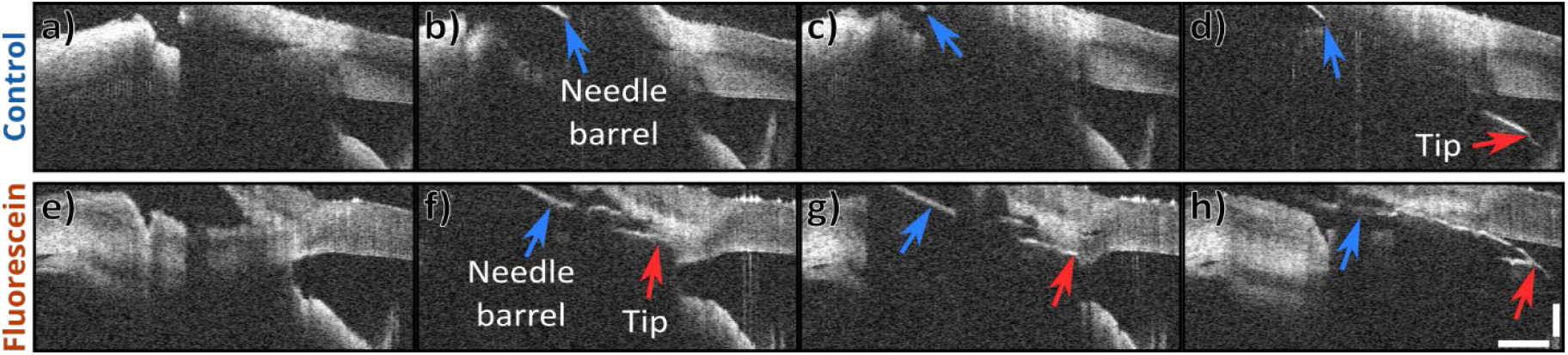
High-speed intrasurgical OCT image series of a trabeculectomy needle insertion maneuver into the anterior chamber of an ex vivo porcine eye. a-d) 7% NaCl solution control eye imaging. a) pre-insertion, b-c) partial insertion, only needle barrel visible (blue arrow), d) fully inserted, needle tip can be visualized beneath the cornea (red arrow). e-h) Corresponding imaging in fluorescein-cleared eye, the needle tip is visible throughout the maneuver (red arrow). 1mm scale.

Here the potential impact of clearing on intrasurgical image guidance is illustrated: in the control case, before insertion, it is difficult to visualize the target AC angle (a). As the needle is inserted to create a drainage channel from the anterior chamber (b, c), it is not possible to visualize the needle tip or target AC angle in the control eye, only the exposed barrel of the needle (blue arrow). Only once the needle is fully inserted into the anterior chamber is the needle tip visible through the cornea (d, red arrow). On the other hand, in the cleared case, the target AC angle is well-visualized pre-insertion (e). As the needle tip is inserted, it remains visible beneath the tissue throughout the entire maneuver into the anterior chamber (f-h). This improved visualization of instrument and target could enable better accuracy in the positioning of the drainage channel, helping to place it sufficiently anterior to avoid the iris, while posterior enough to avoid the cornea.

### 4.3. *Ex vivo* Human Imaging

Following *ex vivo* porcine experiments, cleared imaging in *ex vivo* human eyes was investigated. Analogous to Fig. 3 in the porcine case, Fig. 8 depicts the fluorescein clearing effect in OCT and white-light imaging over time (n=1).

**Figure 8.**
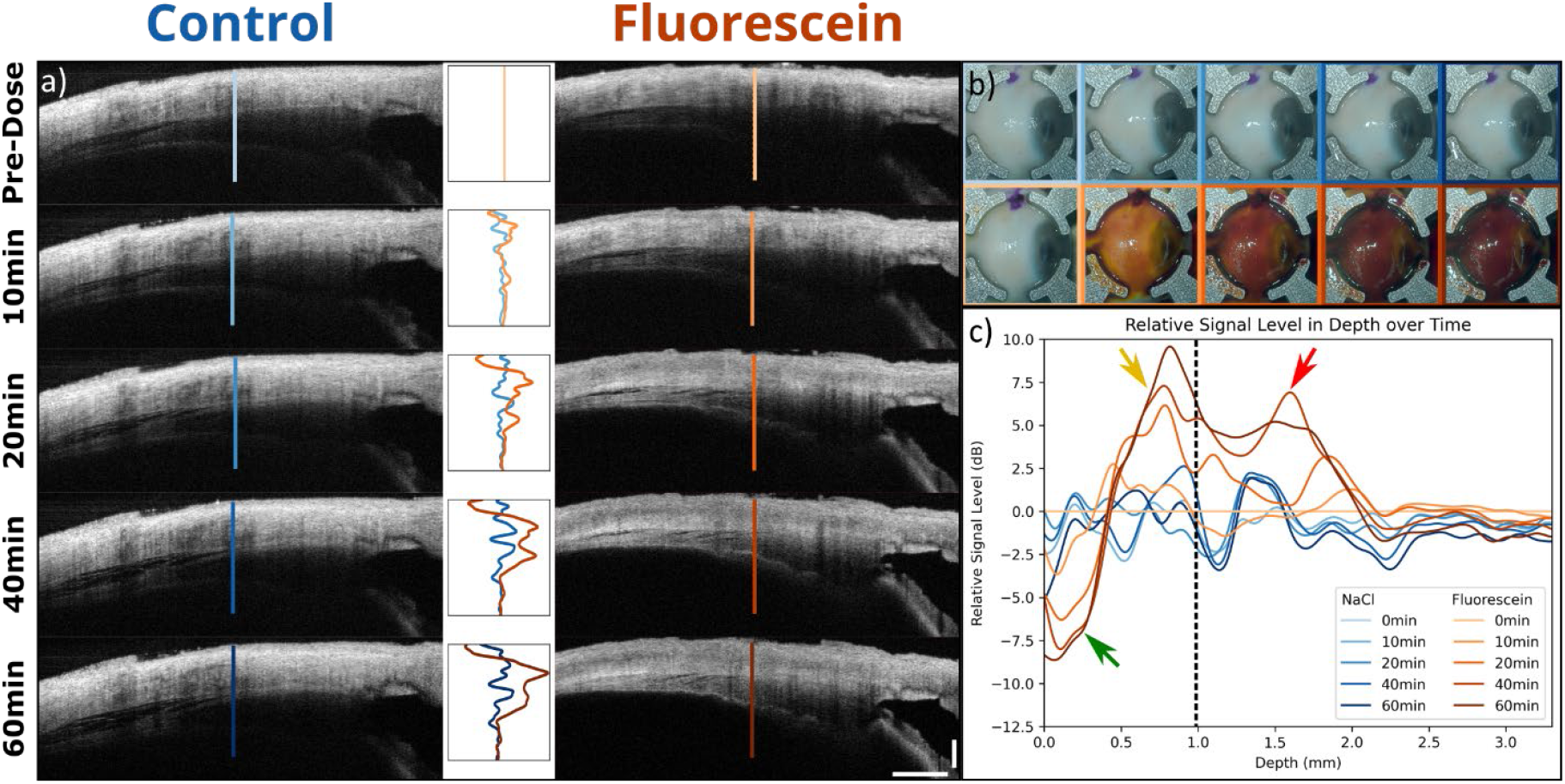
a) Time series OCT imaging of clearing effect over time of salt control solution (left) compared to fluorescein (right) in ex vivo human eyes. Top row is immediately before dosing, followed by 10, 20, 40, and 60 minutes of dose application. Smoothed image signal depth profiles for each image are plotted normalized to the starting profile (middle), with line annotations indicating the position of the plotted a-scan. (b) Representative enface surgical microscopy imaging for salt control (top row) and fluorescein (bottom) over time. (c) Combined plot of all normalized depth profiles over all timepoints, with approximate position of superficial sclera (green arrow), bulk sclera (yellow arrow), choroid-scleral junction (dotted black line), and ciliary body/choroid (red arrow). 1mm scale.

Like the porcine case, a clearing effect over time is apparent when comparing the OCT imaging of salt solution control (8a, left) to the 30% fluorescein treatment (right), as illustrated by the plots of the depth profiles of the relative image signal (middle). Examining the corresponding white-light imaging in 8b, it is noted that the human eye appears to darken to an extent even greater than that observed in the porcine case following treatment. Plotting the depth profiles together in 8c reveals feature similar to those noted in the porcine case (Fig. 3c), with several notable differences. Firstly, the overall length of the profile is shorter due to the lesser thickness of the human sclera. Additionally, the relative gains in the signal of the deeper scleral tissue here (first +10dB peak) are not as great as the porcine eyes following treatment. This is also attributed to the thinner human sclera; with the decreased thickness of the tissue, there is a twofold effect: a greater starting image signal throughout the scleral depth pre-treatment, and there is lesser deep scleral image signal post-treatment due to more complete clearing of the full scleral tissue depth. In combination, this reduces the deep scleral signal increase from clearing in human compared to porcine. However, while this clearing effect does reduce the signal gain in the sclera, this also results in increased signal gain of deeper features below the sclera including choroid/ciliary body (+7.5dB peak, red arrow) compared to porcine.

Like the previous porcine case, a trabeculectomy procedure was performed on a pair of *ex vivo* human control and cleared eyes, with imaging results of the needle insertion step assessed in Fig. 9, analogous to the Fig. 6 porcine results.

**Figure 9.**
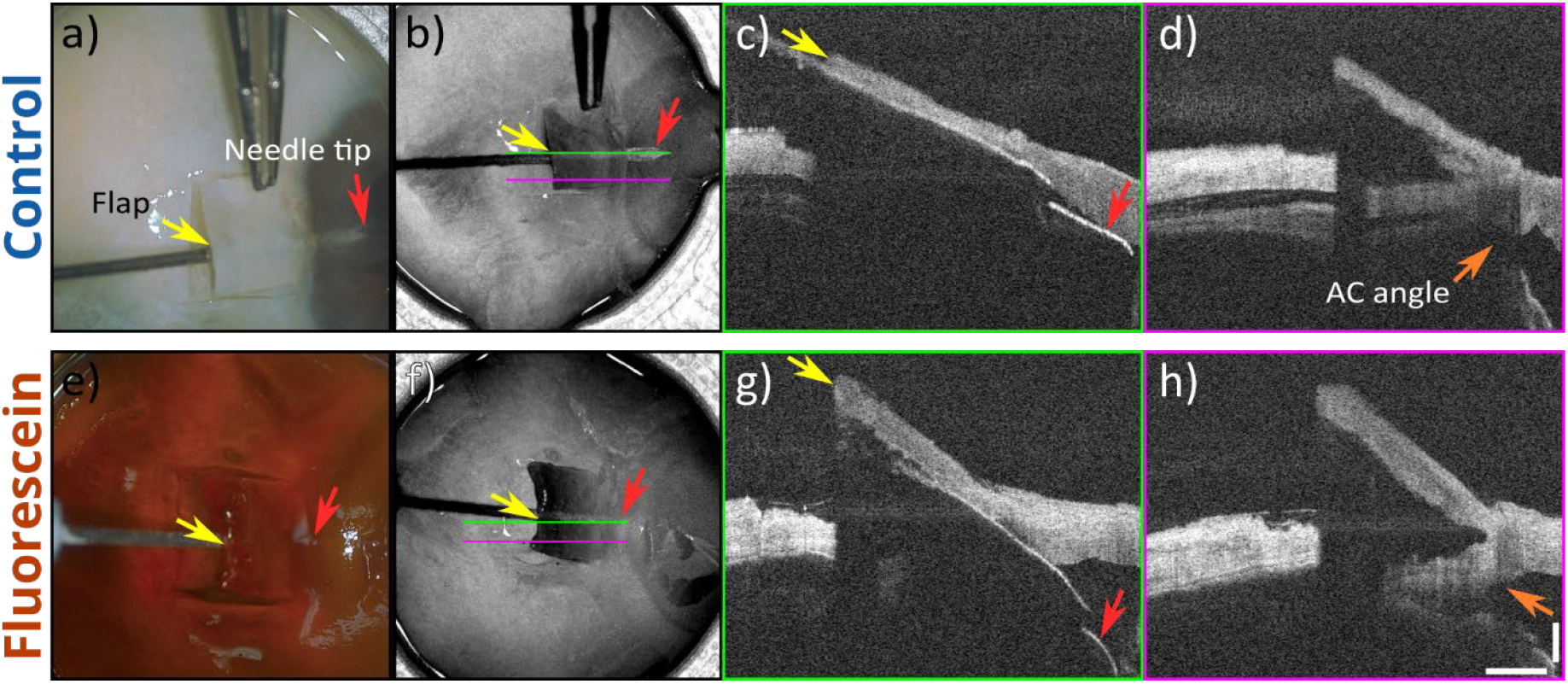
a) Intrasurgical microscopy imaging of ex vivo human eye during trabeculectomy channel creation following 1hr exposure to 7% NaCl solution. Scleral flap (yellow arrow) and needle tip (red) visualized. b) Corresponding OCT enface max-intensity projection. c) B-scan along needle axis (green). d) B-scan adjacent to needle to visualize features previously shadowed by needle, including the flap bed (purple arrow) and AC angle (orange). e-h) Corresponding 30% fluorescein imaging, isoosmolar to 7% NaCl. 1mm scale.

Similar to the porcine case in Fig. 6, clearing improvements to OCT depth visibility are evident. The AC angle visibility increases from control (d, orange arrow) to cleared (h). However, due to the decreased thickness of the flap and sclera in the human case, the visualization gain in the needle is not substantial, with the it clearly visualized beneath the flap in both the control (9c) and treatment (g) cases, although the flap is thinner in the control case. In the control case, a detachment of the choroid can also be seen; this is not an uncommon occurrence in *ex vivo* eye samples in both human and porcine cases (see Fig. 4a, 10% fluorescein for porcine detachment example) and is not considered a result of the applied treatments.

## 5. Discussion

We have demonstrated a novel application of fluorescein dye as a reversible optical clearing agent for increasing OCT imaging depth through highly scattering biological tissue, with potential for applications in the enhancement of intrasurgical OCT image guidance. While transscleral imaging in the context of trabeculectomy was investigated here, this fluorescein tissue clearing could prove useful in several other applications in the anterior segment. As mentioned earlier, enhanced imaging depth would improve glaucoma angle measurements, determination of malignancy margins, verification of implants and injections, and localization of muscle insertions for strabismus surgical treatments. This potential also extends to other challenging image tissue targets beyond ophthalmology, including dermatological, cervical, and neurosurgical imaging. Many applications of OCT in these spaces would benefit from increased depths of imaging for more complete mappings of bulk tissue and vascular network structural and functional information. Similarly, the benefits of clearing are not limited to OCT as a modality; spectroscopy, microscopy, fluoroscopy, endoscopy, photoacoustic, and photodynamic techniques can all potentially fluorescein’s clearing effects^7^. Even within this work, corresponding surgical microscopy imaging suggests that the method can increase the imaging depth of white-light microscopy, as the sclera is observed to darken with clearing as less illumination is backscattered by the sclera and is instead absorbed by the retinal pigmented epithelium (RPE) beneath (Fig. 3b). This effect is even more pronounced in the human case (Fig. 8b), due to the decreased thickness of the human sclera compared to porcine resulting in even greater absorption by RPE the rather than its backscatter by the sclera, darkening the tissue appearance under surgical microscopy. As an accessible, cost-effective, and well-tested agent with an extensive history in clinical applications, fluorescein could serve as a powerful and versatile tool for optical tissue clearing both *in vivo* as in clinical applications, and *ex vivo* in pathological testing and models.

While there is potential for wide applications, there are obvious caveats to discuss. In the future, reversibility and repeatability of the effect in humans should be examined, but this is beyond the scope of initial investigation ascribed to the work here. Regardless, there are still meaningful observations to be made from the human data: although depth of imaging was improved in the human case, the effect was less dramatic compared to porcine; this can be straightforwardly attributed to the lesser thickness of the human sclera; the imaged human sclera has been nearly completely cleared, and thus exhibits less reflectivity than portions of the porcine sclera which have not yet been cleared due to their increased depth from the surface. Combined with the fact that the full thickness of the sclera is initially visible in the human case, more of the visibility change with clearing treatment is seen in the ciliary body deeper below, which exhibits less reflectance than the layers of deep sclera in cleared porcine imaging. It may be that the clearing effect will be more useful in clinical imaging scenarios where the conjunctiva is not removed, further increasing the required depth of imaging, unlike this case. Additionally, fluorescein is typically clinically used at 2% concentration; in this exploratory study to maximize the effect/signal, thus we used a significantly greater concentration of 30%. While this is a relatively high concentration, it is not strictly necessary for it to be this high to achieve a clearing effect. As a study aimed at first characterization of the effect, an excessive upper limit of concentration was applied, but it is expected that lower concentrations will also exhibit noticeable and potentially equivalent clearing given a longer dosing time. Here the effect appears to saturate between 15-30% fluorescein concentration at 1 hour, or 40-60 minutes at 30% concentration. Lesser clearing may even be sufficient for the thinner human sclera. In any case, as this research moves forward, it will be necessary to carefully evaluate the biocompatibility of higher topical concentrations of fluorescein, and consider the feasibility and costs of adding preoperative dosing time to the procedure.

Finally, it should be noted that fluorescein is not the only ophthalmic dye with clearing potential in OCT; another common dye that should be considered in investigations of tissue-clearing applications is indocyanine green (ICG). While it is significantly more costly and is not typically applied topically in ophthalmic applications, it may be spectrally advantaged compared to fluorescein for NIR OCT imaging, with its absorbance peak of ∼700-780nm falling closer to the NIR imaging wavelength of most OCT systems. This may enable increased clearing at lower concentrations and dosing times compared to fluorescein, potentially providing an optical clearing agent with improved biocompatibility and clinical applicability compared to fluorescein.

## Data Availability

Data underlying these results are available upon reasonable request.

## Acknowledgements

Thanks to Michelle McCall for research and administrative support.

